# Novel regulatory and transcriptional networks associated with resistance to platinum-based chemotherapy in ovarian cancer

**DOI:** 10.1101/2020.09.09.289868

**Authors:** Danai G. Topouza, Jihoon Choi, Sean Nesdoly, Anastasiya Tarnouskaya, Christopher J.B. Nicol, Qing Ling Duan

**Affiliations:** Department of Biomedical and Molecular Sciences, Queen’s University, Kingston, Ontario, Canada; School of Computing, Queen’s University, Kingston, Ontario, Canada; Department of Pathology and Molecular Medicine, Queen’s University, Kingston, Ontario, Canada; Division of Cancer Biology and Genetics, Queen’s University Cancer Research Institute, Queen’s University, Kingston, Ontario, Canada

**Author notes:** Correspondence: Qing Ling Duan, Computational Genomics Laboratory, Botterell Hall, Room 530, 18 Stuart St, Kingston, ON, Canada, K7L 3N6.

**Keywords:** Drug Resistance, Neoplasm, Carcinoma, Ovarian Epithelial, High-Throughput RNA Sequencing, Gene Regulatory Networks, MicroRNAs, Quantitative Trait Loci

## Abstract

**Background:** High-grade serous ovarian cancer (HGSOC) is a highly lethal gynecologic cancer, in part due to resistance to platinum-based chemotherapy reported among 20% of patients. This study aims to elucidate the biological mechanisms underlying chemotherapy resistance, which remain poorly understood.

**Methods:** Sequencing data (mRNA and microRNA) from HGSOC patients were analyzed to identify differentially expressed genes and co-expressed transcript networks associated with chemotherapy response. Initial analyses used datasets from The Cancer Genome Atlas and then replicated in two independent cancer cohorts. Moreover, transcript expression datasets and genomics data (i.e. single nucleotide polymorphisms) were integrated to determine potential regulation of the associated mRNA networks by microRNAs and expression quantitative trait loci (eQTLs).

**Results:** In total, 196 differentially expressed mRNAs were enriched for adaptive immunity and translation, and 21 differentially expressed microRNAs were associated with angiogenesis. Moreover, co-expression network analysis identified two mRNA networks associated with chemotherapy response, which were enriched for ubiquitination and lipid metabolism, as well as three associated microRNA networks enriched for lipoprotein transport and oncogenic pathways. In addition, integrative analyses revealed potential regulation of the mRNA networks by the associated microRNAs and eQTLs.

**Conclusion:** We report novel transcriptional networks and pathways associated with resistance to platinum-based chemotherapy among HGSOC patients. These results aid our understanding of the effector networks and regulators of chemotherapy response, which will improve drug efficacy and identify novel therapeutic targets for ovarian cancer.

## 1. Introduction

High-grade serous ovarian cancer (HGSOC) is a highly lethal gynecologic cancer, in part due to resistance to first-line, platinum-based chemotherapy treatment among 20% of patients [1]. Chemotherapy resistant patients have a significantly shorter overall survival (OS) than sensitive patients, and many experience tumor recurrence within six months of completing chemotherapy [2]. There is currently no strategy for predicting response to platinum-based chemotherapy, which reflects our limited understanding of the underlying molecular mechanisms of chemoresistance [3].

Earlier studies reporting gene expression signatures associated with chemoresistance in ovarian cancer patients had used univariate analysis methods, which assume that chemotherapy response is driven by a single gene [4]. However, it is well established that chemotherapy response, like other drug response outcomes, is a complex multifactorial trait modulated by multiple genes contributing to one or more biological pathways [5].

In this study, we apply both univariate and multivariate analysis methods to provide the first evidence of novel mRNA and microRNA (miRNA) signalling pathways and networks associated with chemotherapy response in HGSOC patients. We further unveil miRNAs and expression quantitative trait loci (eQTLs) correlated with the expression of significant transcripts. These findings are validated using two independent cancer cohorts and provide important therapeutic strategies for regulating platinum-based chemotherapy response in HGSOC patients.

## 2. Materials and Methods

### 2.1 Chemotherapy response classification

Sequencing of mRNA and miRNA was derived from chemotherapy-naïve tumors of 191 and 205 HGSOC patients of TCGA, respectively [6]. Patients who received platinum-based adjuvant chemotherapy were selected and classified for chemotherapy response based on their platinum-free interval. Sensitive patients remained cancer-free for at least 12 months after chemotherapy completion, whereas resistant patients experienced cancer recurrence within 6 months (**Table 1**; **Supplementary Methods**).

**Table 1.**
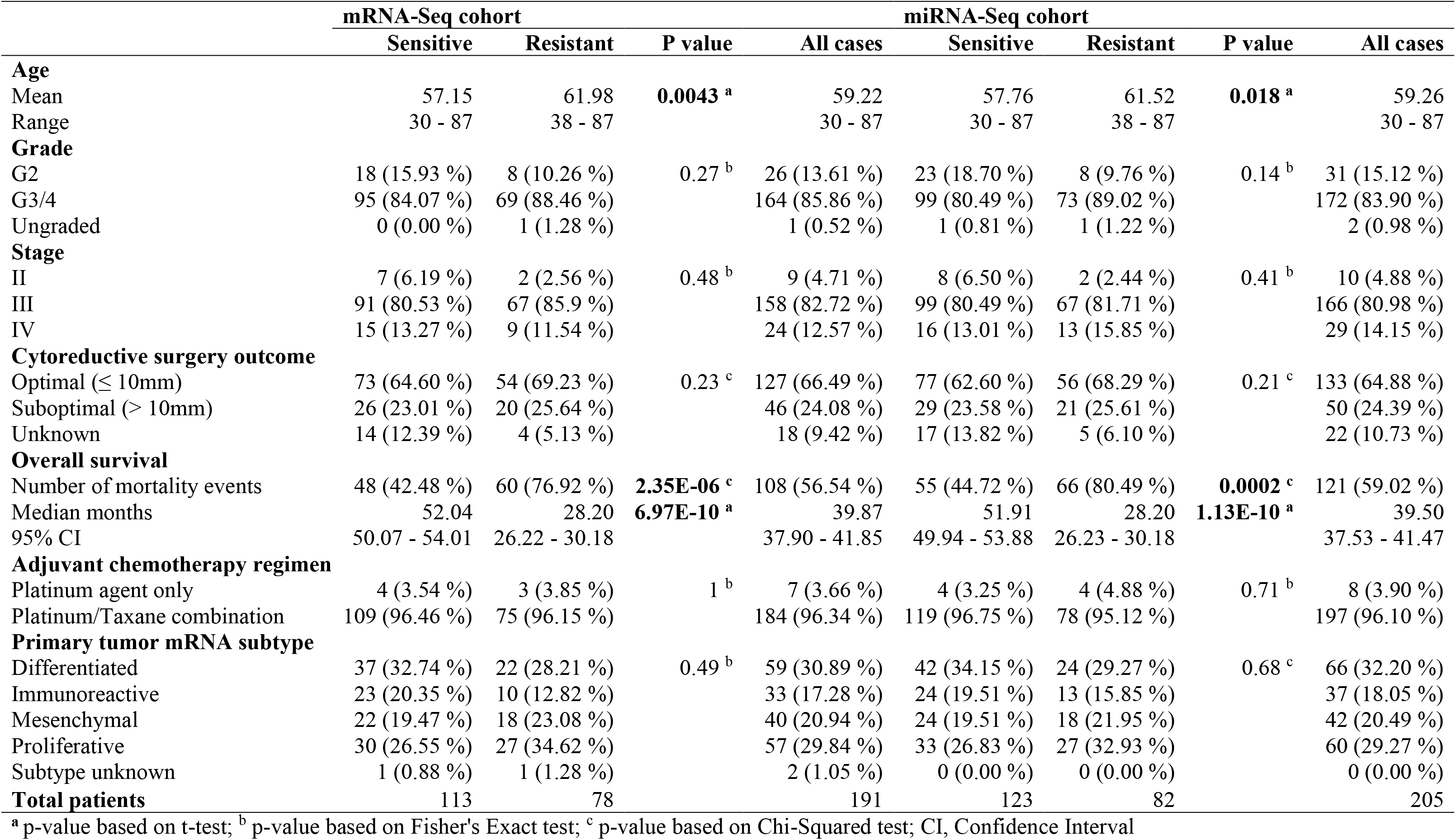
Characteristics of the HGSOC patient cohorts with mRNA-Seq and miRNA-Seq data from TCGA.

|  | mRNA-Seq cohort |  |  | miRNA-Seq cohort |  |  |  |  |
| --- | --- | --- | --- | --- | --- | --- | --- | --- |
|  | Sensitive | Resistant | P value | All cases | Sensitive | Resistant | P value | All cases |
| **Age** |  |  |  |  |  |  |  |  |
| Mean | 57.15 | 61.98 | **0.0043<sup>a</sup>** | 59.22 | 57.76 | 61.52 | **0.018<sup>a</sup>** | 59.26 |
| Range | 30 - 87 | 38 - 87 |  | 30 - 87 | 30 - 87 | 38 - 87 |  | 30 - 87 |
| **Grade** |  |  |  |  |  |  |  |  |
| G2 | 18 (15.93 %) | 8 (10.26 %) | 0.27<sup>b</sup> | 26 (13.61 %) | 23 (18.70 %) | 8 (9.76 %) | 0.14<sup>b</sup> | 31 (15.12 %) |
| G3/4 | 95 (84.07 %) | 69 (88.46 %) |  | 164 (85.86 %) | 99 (80.49 %) | 73 (89.02 %) |  | 172 (83.90 %) |
| Ungraded | 0 (0.00 %) | 1 (1.28 %) |  | 1 (0.52 %) | 1 (0.81 %) | 1 (1.22 %) |  | 2 (0.98 %) |
| **Stage** |  |  |  |  |  |  |  |  |
| II | 7 (6.19 %) | 2 (2.56 %) | 0.48<sup>b</sup> | 9 (4.71 %) | 8 (6.50 %) | 2 (2.44 %) | 0.41<sup>b</sup> | 10 (4.88 %) |
| III | 91 (80.53 %) | 67 (85.9 %) |  | 158 (82.72 %) | 99 (80.49 %) | 67 (81.71 %) |  | 166 (80.98 %) |
| IV | 15 (13.27 %) | 9 (11.54 %) |  | 24 (12.57 %) | 16 (13.01 %) | 13 (15.85 %) |  | 29 (14.15 %) |
| **Cytoreductive surgery outcome** |  |  |  |  |  |  |  |  |
| Optimal ($\leq 10$mm) | 73 (64.60 %) | 54 (69.23 %) | 0.23<sup>c</sup> | 127 (66.49 %) | 77 (62.60 %) | 56 (68.29 %) | 0.21<sup>c</sup> | 133 (64.88 %) |
| Suboptimal ($> 10$mm) | 26 (23.01 %) | 20 (25.64 %) | | 46 (24.08 %) | 29 (23.58 %) | 21 (25.61 %) | | 50 (24.39 %) |
| Unknown | 14 (12.39 %) | 4 (5.13 %) |  | 18 (9.42 %) | 17 (13.82 %) | 5 (6.10 %) |  | 22 (10.73 %) |
| **Overall survival** |  |  |  |  |  |  |  |  |
| Number of mortality events | 48 (42.48 %) | 60 (76.92 %) | **2.35E-06<sup>c</sup>** | 108 (56.54 %) | 55 (44.72 %) | 66 (80.49 %) | **0.0002<sup>c</sup>** | 121 (59.02 %) |
| Median months | 52.04 | 28.20 | **6.97E-10<sup>a</sup>** | 39.87 | 51.91 | 28.20 | **1.13E-10<sup>a</sup>** | 39.50 |
| 95% CI | 50.07 - 54.01 | 26.22 - 30.18 |  | 37.90 - 41.85 | 49.94 - 53.88 | 26.23 - 30.18 |  | 37.53 - 41.47 |
| **Adjuvant chemotherapy regimen** |  |  |  |  |  |  |  |  |
| Platinum agent only | 4 (3.54 %) | 3 (3.85 %) | 1<sup>b</sup> | 7 (3.66 %) | 4 (3.25 %) | 4 (4.88 %) | 0.71<sup>b</sup> | 8 (3.90 %) |
| Platinum/Taxane combination | 109 (96.46 %) | 75 (96.15 %) |  | 184 (96.34 %) | 119 (96.75 %) | 78 (95.12 %) |  | 197 (96.10 %) |
| **Primary tumor mRNA subtype** |  |  |  |  |  |  |  |  |
| Differentiated | 37 (32.74 %) | 22 (28.21 %) | 0.49<sup>b</sup> | 59 (30.89 %) | 42 (34.15 %) | 24 (29.27 %) | 0.68<sup>c</sup> | 66 (32.20 %) |
| Immunoreactive | 23 (20.35 %) | 10 (12.82 %) |  | 33 (17.28 %) | 24 (19.51 %) | 13 (15.85 %) |  | 37 (18.05 %) |
| Mesenchymal | 22 (19.47 %) | 18 (23.08 %) |  | 40 (20.94 %) | 24 (19.51 %) | 18 (21.95 %) |  | 42 (20.49 %) |
| Proliferative | 30 (26.55 %) | 27 (34.62 %) |  | 57 (29.84 %) | 33 (26.83 %) | 27 (32.93 %) |  | 60 (29.27 %) |
| Subtype unknown | 1 (0.88 %) | 1 (1.28 %) |  | 2 (1.05 %) | 0 (0.00 %) | 0 (0.00 %) |  | 0 (0.00 %) |
| **Total patients** | 113 | 78 |  | 191 | 123 | 82 |  | 205 |

<sup>a</sup> p-value based on t-test; <sup>b</sup> p-value based on Fisher's Exact test; <sup>c</sup> p-value based on Chi-Squared test; CI, Confidence Interval

### 2.2 Processing of sequencing data

Sequencing reads from mRNA were downloaded as FASTQ files (i.e. level 1 data from TCGA), filtered for base-quality, aligned, and quantified (detailed in **Supplementary Methods)**. Sequencing of miRNA data were downloaded as quantified expression files (i.e. level 3 data from TCGA). Both mRNA and miRNA datasets underwent outlier detection, normalization, and non-specific filtering, resulting in 49,116 mRNA and 4,479 miRNA transcripts for further analyses.

### 2.3 Differential expression analysis

Differentially expressed mRNA and miRNA transcripts were detected between sensitive and resistant patients using a negative binomial generalized linear model (GLM) in the DESeq2 R package [7]. This analysis controlled for patients’ ages at diagnosis, as resistant patients were significantly older (**Table 1**). The Benjamini-Hochberg method corrected for multiple testing.

### 2.4 Weighted co-expression network analysis

The weighted gene co-expression network analysis (WGCNA) R package [8] was used to identify modules of co-expressed mRNA and miRNA transcripts using an unsupervised machine learning approach. The first principal component of expression values was calculated for each transcript module, resulting in an eigengene value. Module eigengenes were used to determine association with chemotherapy response using a GLM, adjusted for patients’ age as a covariate (**Supplementary Methods**).

### 2.5 Pathway enrichment analysis

Pathway enrichment analysis was used to determine over-representation of biological pathways from lists of differentially expressed transcripts (mRNA and miRNAs) and co-expression networks (**Supplementary Methods**).

### 2.6 eQTL analysis

Germline single nucleotide polymorphisms (SNPs) from TCGA-HGSOC patients were imputed as described by Choi *et al.* [9] before undergoing quality control and linkage disequilibrium-based pruning, retaining 1,722,608 common SNPs for analysis. SNPs were integrated with patient mRNA-seq data (n=167) and miRNA-seq data (n=178) to identify correlations with transcript expression (eQTLs) using the MatrixEQTL R package [10] (**Supplementary Methods**).

### 2.7 mRNA-microRNA integration

Potential regulation of mRNA networks by miRNAs was tested using data from 165 patients by applying the Spearman correlation of module eigengenes from the mRNA and miRNA co-expression networks. Results were validated using miRNet [11], a database of experimentally validated mRNA-miRNA interactions, and miRGate [12], a database of predicted mRNA-miRNA interactions based on ribonucleotide motif matching.

### 2.8 Replication cohort and analysis

Replication of differentially expressed transcripts used the Kaplan-Meier analysis tool KM Plotter [13] to test the association of each transcript with progression-free survival (PFS) or OS in two independent cancer cohorts. First, mRNA transcripts were replicated with PFS from the Australian Ovarian Cancer Study (AOCS; GSE9891; n=285) [14]. Next, miRNA isoforms were replicated with OS in the TCGA bladder urothelial carcinoma cohort (BLCA) (n=409) [15]. Validation of transcript networks used the Prognostic Index estimation method [16] using data from these two cohorts (**Supplementary Methods**).

## 3. Results

Differential expression analysis identified 196 mRNAs associated with chemotherapy response (adjusted p<0.05) that map to 190 unique genes (**Figure 1A, Table S1**). Pathway enrichment analysis of these associated transcripts indicated enrichment of 41 annotation terms, including B-cell receptor regulation, complement activation, and peptide chain elongation (**Table 2**).

**Table 2.**
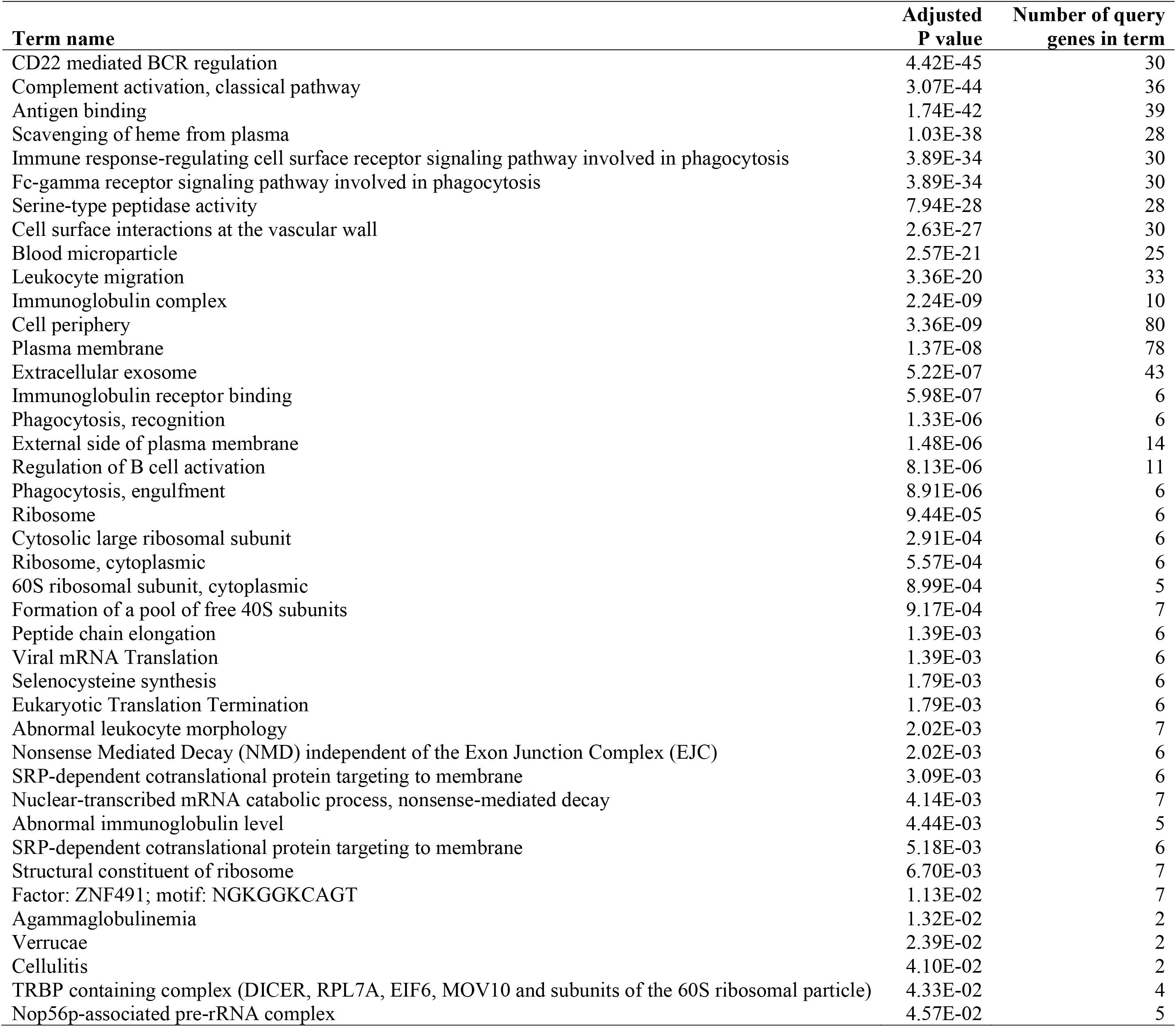
Significantly enriched annotation terms for differentially expressed mRNA transcripts.

| **Term name** | **Adjusted**<br>**P value** | **Number of query**<br>**genes in term** |
| --- | --- | --- |
| CD22 mediated BCR regulation | 4.42E-45 | 30 |
| Complement activation, classical pathway | 3.07E-44 | 36 |
| Antigen binding | 1.74E-42 | 39 |
| Scavenging of heme from plasma | 1.03E-38 | 28 |
| Immune response-regulating cell surface receptor signaling pathway involved in phagocytosis | 3.89E-34 | 30 |
| Fc-gamma receptor signaling pathway involved in phagocytosis | 3.89E-34 | 30 |
| Serine-type peptidase activity | 7.94E-28 | 28 |
| Cell surface interactions at the vascular wall | 2.63E-27 | 30 |
| Blood microparticle | 2.57E-21 | 25 |
| Leukocyte migration | 3.36E-20 | 33 |
| Immunoglobulin complex | 2.24E-09 | 10 |
| Cell periphery | 3.36E-09 | 80 |
| Plasma membrane | 1.37E-08 | 78 |
| Extracellular exosome | 5.22E-07 | 43 |
| Immunoglobulin receptor binding | 5.98E-07 | 6 |
| Phagocytosis, recognition | 1.33E-06 | 6 |
| External side of plasma membrane | 1.48E-06 | 14 |
| Regulation of B cell activation | 8.13E-06 | 11 |
| Phagocytosis, engulfment | 8.91E-06 | 6 |
| Ribosome | 9.44E-05 | 6 |
| Cytosolic large ribosomal subunit | 2.91E-04 | 6 |
| Ribosome, cytoplasmic | 5.57E-04 | 6 |
| 60S ribosomal subunit, cytoplasmic | 8.99E-04 | 5 |
| Formation of a pool of free 40S subunits | 9.17E-04 | 7 |
| Peptide chain elongation | 1.39E-03 | 6 |
| Viral mRNA Translation | 1.39E-03 | 6 |
| Selenocysteine synthesis | 1.79E-03 | 6 |
| Eukaryotic Translation Termination | 1.79E-03 | 6 |
| Abnormal leukocyte morphology | 2.02E-03 | 7 |
| Nonsense Mediated Decay (NMD) independent of the Exon Junction Complex (EJC) | 2.02E-03 | 6 |
| SRP-dependent cotranslational protein targeting to membrane | 3.09E-03 | 6 |
| Nuclear-transcribed mRNA catabolic process, nonsense-mediated decay | 4.14E-03 | 7 |
| Abnormal immunoglobulin level | 4.44E-03 | 5 |
| SRP-dependent cotranslational protein targeting to membrane | 5.18E-03 | 6 |
| Structural constituent of ribosome | 6.70E-03 | 7 |
| Factor: ZNF491; motif: NGKGGKCAGT | 1.13E-02 | 7 |
| Agammaglobulinemia | 1.32E-02 | 2 |
| Verrucae | 2.39E-02 | 2 |
| Cellulitis | 4.10E-02 | 2 |
| TRBP containing complex (DICER, RPL7A, EIF6, MOV10 and subunits of the 60S ribosomal particle) | 4.33E-02 | 4 |
| Nop56p-associated pre-rRNA complex | 4.57E-02 | 5 |

**Figure 1.**
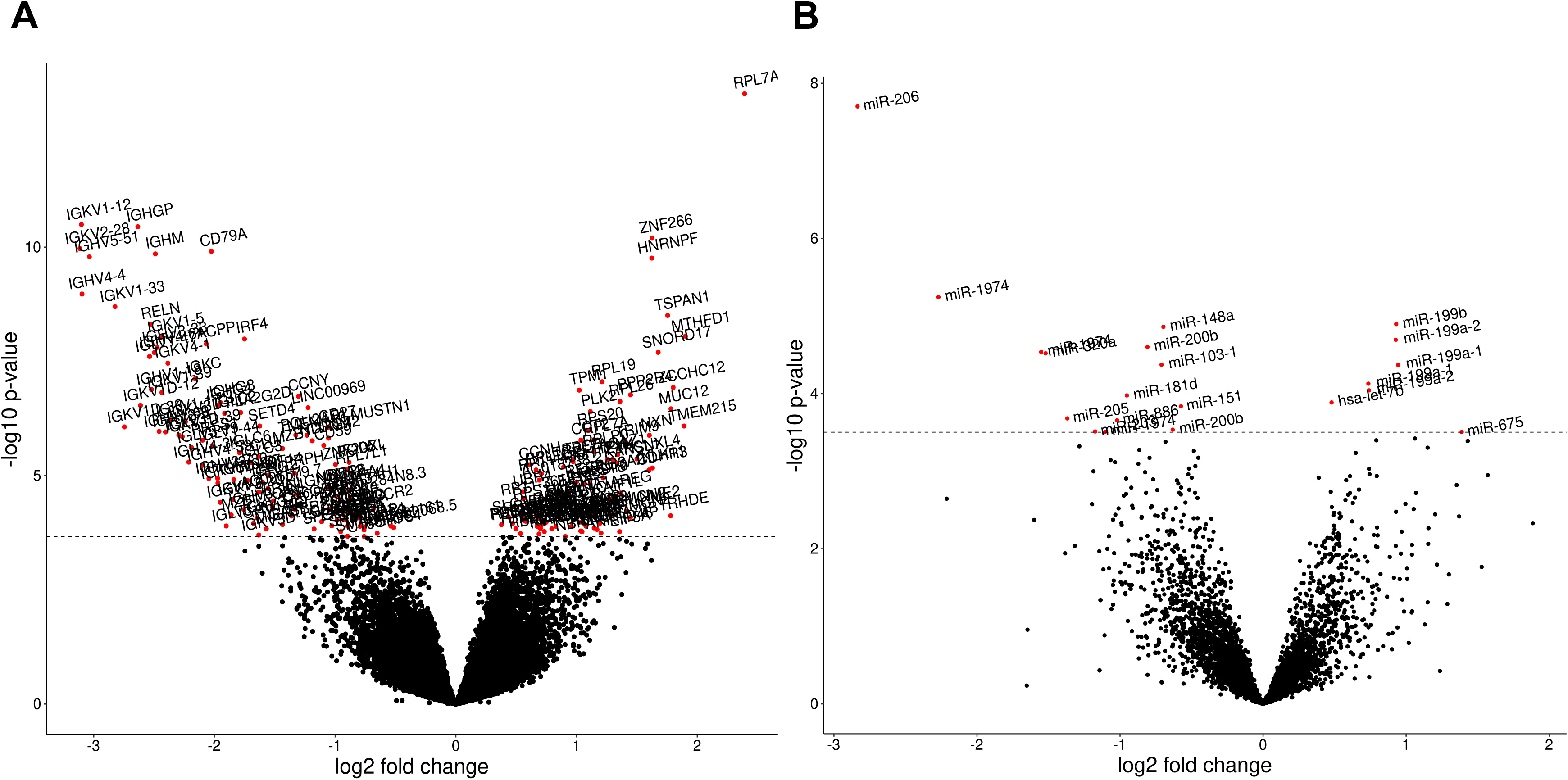
Differential expression analysis of mRNA and miRNA transcripts. (A) Significant differentially expressed mRNA transcripts (n=196) are shown in red. The Benjamini-Hochberg adjusted significance threshold is indicated by the dashed line. Transcripts with a positive fold change are upregulated in resistant patients, whereas transcripts with a negative fold change are downregulated in resistant patients. (B) Significant differentially expressed miRNA isoforms (n=21) are shown in red. The Benjamini-Hochberg adjusted significance threshold is indicated by the dashed line. Transcripts with a positive fold change are upregulated in resistant patients, whereas transcripts with a negative fold change are downregulated in resistant patients.

Differential miRNA expression analysis revealed 21 differentially expressed miRNA isoforms (adjusted p<0.05), which map to 16 unique miRNAs (**Figure 1B, Table S2**). Pathway enrichment analysis of these miRNA isoforms revealed 16 pathways, such as blood vessel morphogenesis and negative regulation of autophagy (**Table 3**).

**Table 3.**
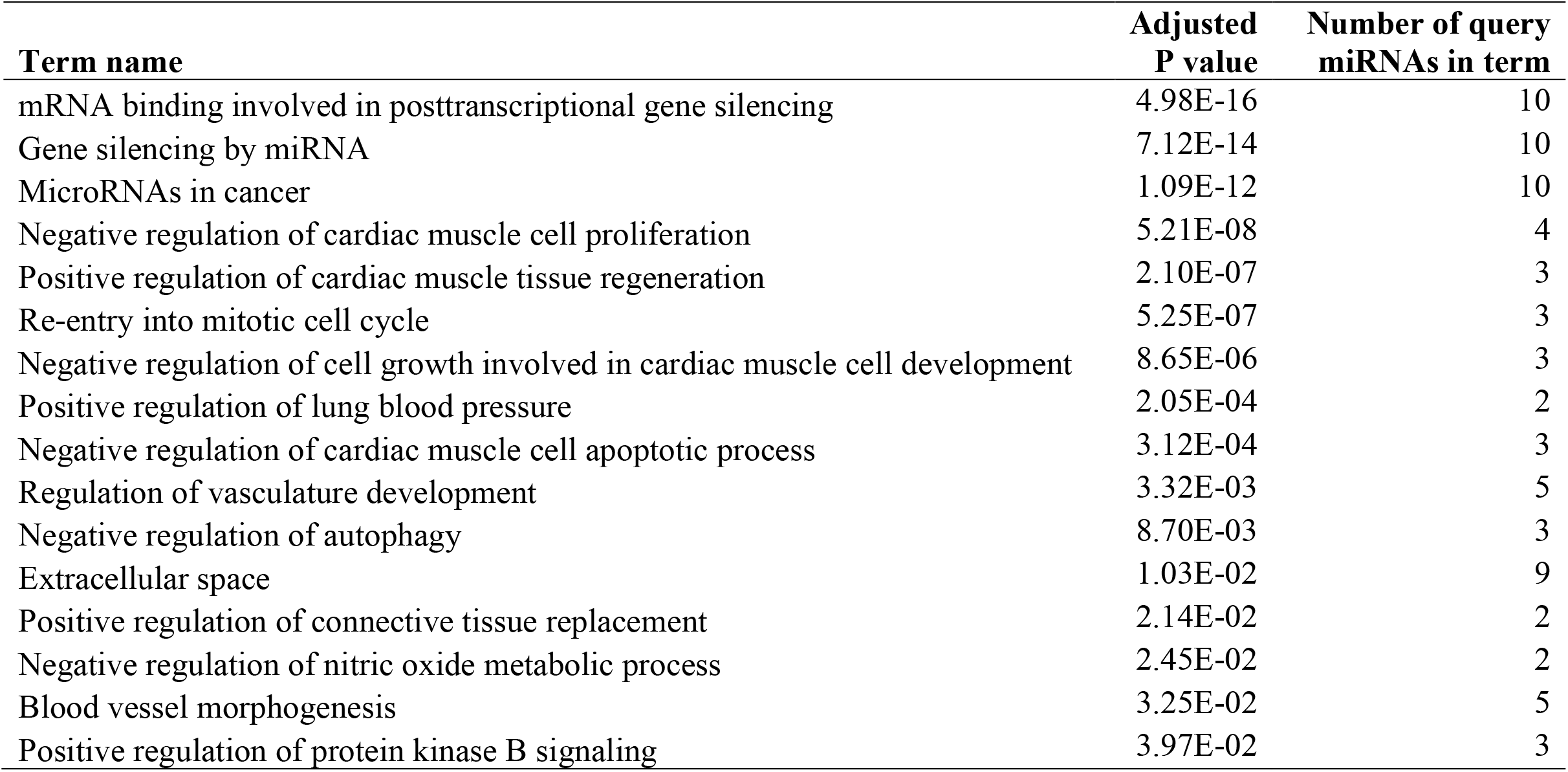
Significantly enriched annotation terms for differentially expressed miRNA isoforms.

| **Term name** | **Adjusted**<br>**P value** | **Number of query**<br>**miRNAs in term** |
| --- | --- | --- |
| mRNA binding involved in posttranscriptional gene silencing | 4.98E-16 | 10 |
| Gene silencing by miRNA | 7.12E-14 | 10 |
| MicroRNAs in cancer | 1.09E-12 | 10 |
| Negative regulation of cardiac muscle cell proliferation | 5.21E-08 | 4 |
| Positive regulation of cardiac muscle tissue regeneration | 2.10E-07 | 3 |
| Re-entry into mitotic cell cycle | 5.25E-07 | 3 |
| Negative regulation of cell growth involved in cardiac muscle cell development | 8.65E-06 | 3 |
| Positive regulation of lung blood pressure | 2.05E-04 | 2 |
| Negative regulation of cardiac muscle cell apoptotic process | 3.12E-04 | 3 |
| Regulation of vasculature development | 3.32E-03 | 5 |
| Negative regulation of autophagy | 8.70E-03 | 3 |
| Extracellular space | 1.03E-02 | 9 |
| Positive regulation of connective tissue replacement | 2.14E-02 | 2 |
| Negative regulation of nitric oxide metabolic process | 2.45E-02 | 2 |
| Blood vessel morphogenesis | 3.25E-02 | 5 |
| Positive regulation of protein kinase B signaling | 3.97E-02 | 3 |

WGCNA of the mRNA transcripts resulted in 58 co-expression modules (**Table S3**), of which two were associated with platinum-based chemotherapy. First, the *lavenderblush3* module is negatively associated with chemotherapy resistance (p-value=0.016; **Figure 2A**). This module contains 39 transcripts, mapping to 31 unique genes (**Figure 2B**). Pathway analysis indicates enrichment of biological pathways related to protein ubiquitination, and the binding motif for transcription factor GABP-alpha (**Table 4**). Second, the *darkolivegreen* module is significantly upregulated in chemo-resistant patients (p-value=0.032) (**Figure 2A**) and contains 82 transcripts mapping to 80 unique genes (**Figure 2B**). This module is significantly enriched for the protein-containing complex term, as well as for 8 pathways involved in fatty acid metabolism with nominal significance (**Table 4**).

**Table 4.**
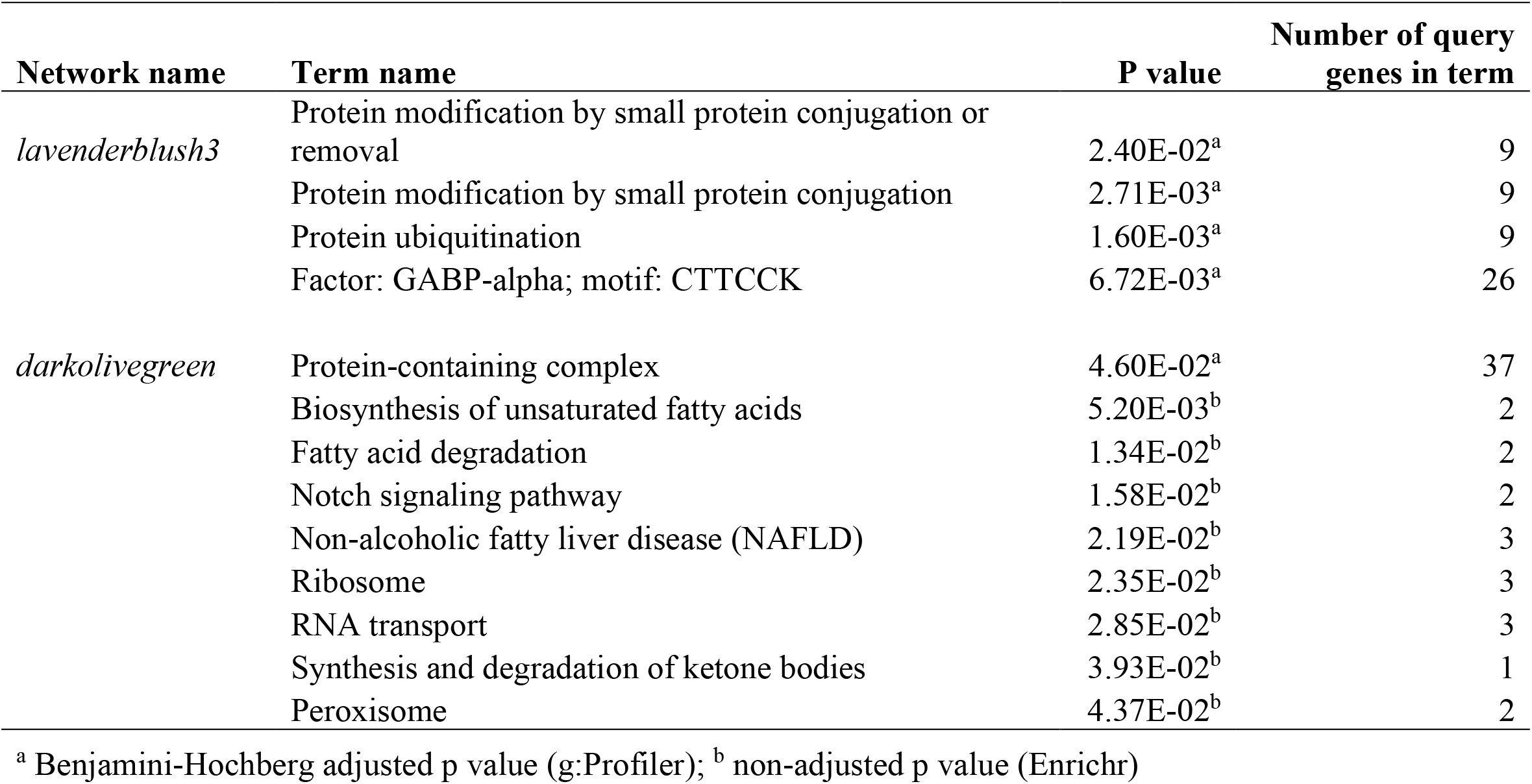
Significantly enriched terms in significant mRNA transcript co-expression networks.

**Figure 2.**
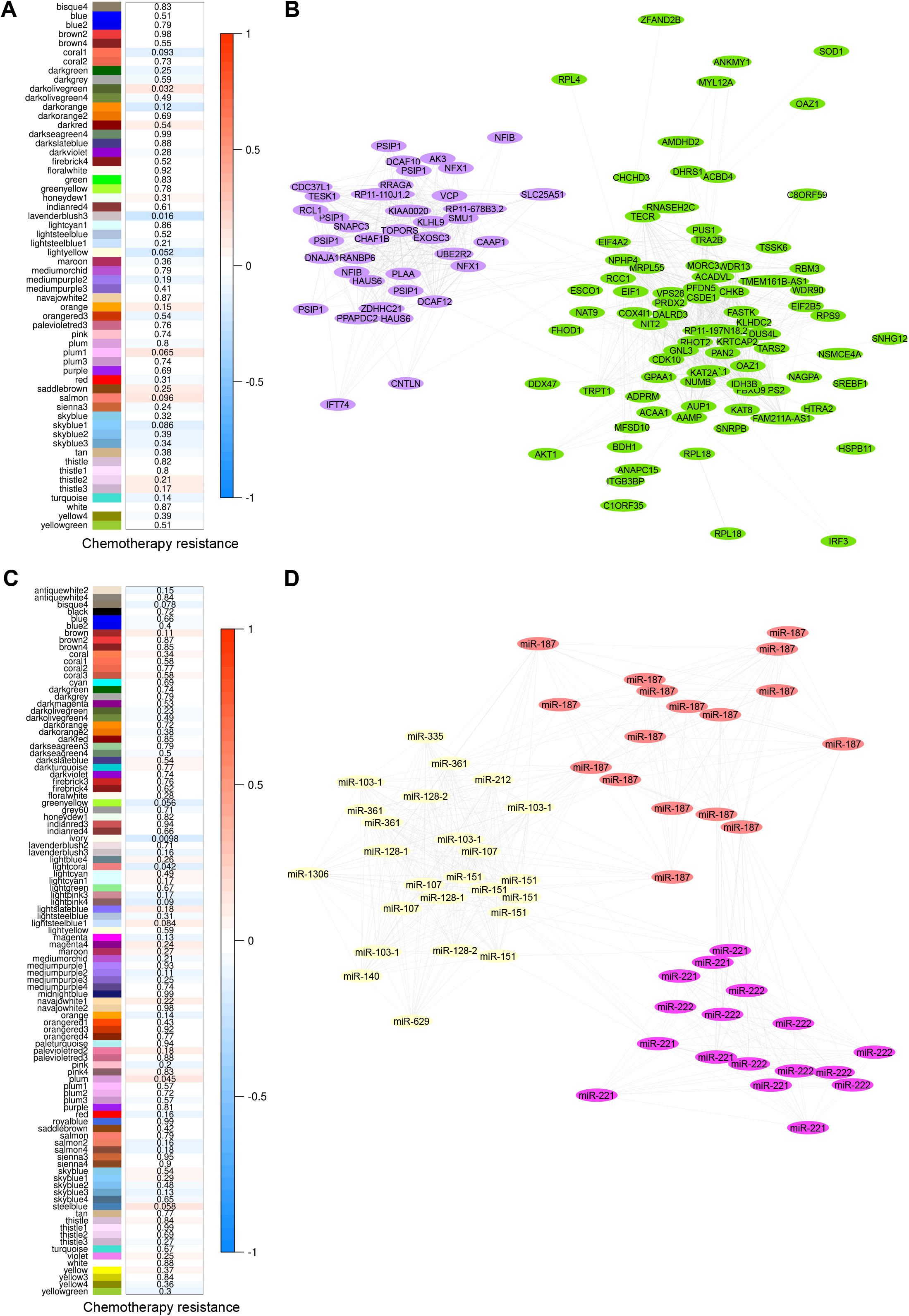
Weighted transcript co-expression network analysis of mRNA and miRNA transcripts. (A) Heatmap of the correlation between the 58 mRNA co-expression modules with platinum-based chemotherapy resistance, red indicating a positive effect and blue a negative effect. P-values are indicated for each module. Two mRNA co-expression modules are significantly associated with chemotherapy resistance (*lavenderblush3*, *darkolivegreen*). (B) The mRNA modules *lavenderblush3* and *darkolivegreen* are visualized in their respective colors. Each node represents a transcript, and each edge represents a connection or co-expression. The distance between nodes is the connection weight, where more similar transcripts are plotted closer. (C) Heatmap of the correlation between the 100 miRNA modules with platinum-based chemotherapy resistance, red indicating a positive effect and blue a negative effect. P-values are indicated for each module. Three miRNA co-expression modules are significantly associated with chemotherapy resistance (*ivory*, *lightcoral*, and *plum*). (D) The miRNA modules *lightcoral*, *plum*, and *ivory* are visualized in their respective colors. Each node represents one miRNA isoform, and each edge represents a connection or co-expression. The distance between nodes is the connection weight, where more similar transcripts are plotted closer.

WGCNA analysis of the miRNA dataset constructed 100 co-expression modules (**Table S4**), of which three are associated with chemotherapy response. The *ivory* (p-value=0.0097) and *lightcoral* (p-value=0.042) modules are negatively associated with chemotherapy resistance, while the third *plum* network (p-value = 0.045) is positively associated with chemotherapy resistance (**Figure 2C**). The *ivory* network consisted of 25 miRNA isoforms mapping to 11 unique miRNAs, which are enriched for 7 pathways and functions, including regulation of lipoprotein transport and cholesterol efflux (**Table 5**). The *lightcoral* module consists of 17 isoforms of miR-187 and the *plum* network consists of 17 isoforms of miR-221 and miR-222 (**Figure 2D**). While no pathway annotations were derived for the *lighcoral* module, the *plum* module is enriched for 18 pathways and oncogenic functions, such as inhibition of the TRAIL-activated apoptotic pathway and inflammatory cytokine production, and up-regulation of protein kinase B signaling (**Table 5**).

**Table 5.** Significantly enriched terms in significant miRNA isoform co-expression networks.

| Network name | Term name | Adjusted P value | Number of query miRNAs in term |
| --- | --- | --- | --- |
| *ivory* | mRNA binding involved in posttranscriptional gene silencing | 1.16E-09 | 7 |
|  | Gene silencing by miRNA | 3.61E-08 | 7 |
|  | Extracellular space | 4.24E-05 | 10 |
|  | MicroRNAs in cancer | 9.55E-05 | 5 |
|  | Regulation of lipoprotein transport | 1.22E-03 | 2 |
|  | Negative regulation of cholesterol efflux | 2.69E-02 | 2 |
|  | Negative regulation of lipoprotein particle clearance | 4.27E-02 | 2 |
| *plum* | Positive regulation of Schwann cell proliferation involved in axon regeneration | 2.55E-06 | 2 |
|  | Negative regulation of hematopoietic stem cell proliferation | 2.55E-06 | 2 |
|  | Positive regulation of Schwann cell migration | 7.66E-06 | 2 |
|  | Negative regulation of TRAIL-activated apoptotic signaling pathway | 7.66E-06 | 2 |
|  | Negative regulation of cell adhesion molecule production | 1.53E-05 | 2 |
|  | Negative regulation by host of viral genome replication | 5.36E-05 | 2 |
|  | Negative regulation of leukocyte adhesion to vascular endothelial cell | 1.40E-04 | 2 |
|  | Negative regulation of cytokine production involved in inflammatory response | 2.32E-04 | 2 |
|  | Positive regulation of erythrocyte differentiation | 1.19E-03 | 2 |
|  | Positive regulation of G1/S transition of mitotic cell cycle | 1.61E-03 | 2 |
|  | Positive regulation of vascular smooth muscle cell proliferation | 2.53E-03 | 2 |
|  | Positive regulation of epithelial to mesenchymal transition | 3.00E-03 | 2 |
|  | Negative regulation of muscle cell differentiation | 5.15E-03 | 2 |
|  | Negative regulation of translation, ncRNA-mediated | 6.17E-03 | 2 |
|  | Negative regulation of sprouting angiogenesis | 8.07E-03 | 2 |
|  | Positive regulation of epithelial cell migration | 3.33E-02 | 2 |
|  | Positive regulation of protein kinase B signaling | 3.58E-02 | 2 |
|  | mRNA binding involved in posttranscriptional gene silencing | 4.25E-02 | 2 |

Integrative analysis with germline SNP data identified 268 unique cis-eQTLs associated with the expression of mRNAs and miRNAs which were correlated with chemotherapy response in differential expression and network analysis. A total of 248 SNPs are associated with the expression of 55 significant mRNAs, and 20 SNPs are associated with the expression of 7 significant miRNAs (**Table S5**). Of the 268 eQTLs, 118 are novel whereas 126 are previously known, and 24 are not yet recorded in the annotation database. The majority (227) are predicted to alter regulatory motifs, and 67 are associated with 94 human phenotypes from published genome-wide association studies. The most common phenotypes are related to triglycerides, high-density lipoprotein (HDL), and low-density lipoprotein (LDL) cholesterol.

Integration of the associated mRNA and miRNA networks determine that the *plum* miRNA network significantly correlates with the *lavenderblush3* mRNA network (Spearman’s ρ = −0.26, p < 0.001), and the *ivory* miRNA network significantly correlates with the *darkolivegreen* mRNA network (Spearman’s ρ = −0.17, p = 0.023). Annotations using miRNet and miRGate determined that 20 of these mRNA-miRNA interactions are experimentally validated, while 15 others are supported by *in silico* predictions (**Table 6, Figure 3, Supplementary Data 1**).

**Table 6.**
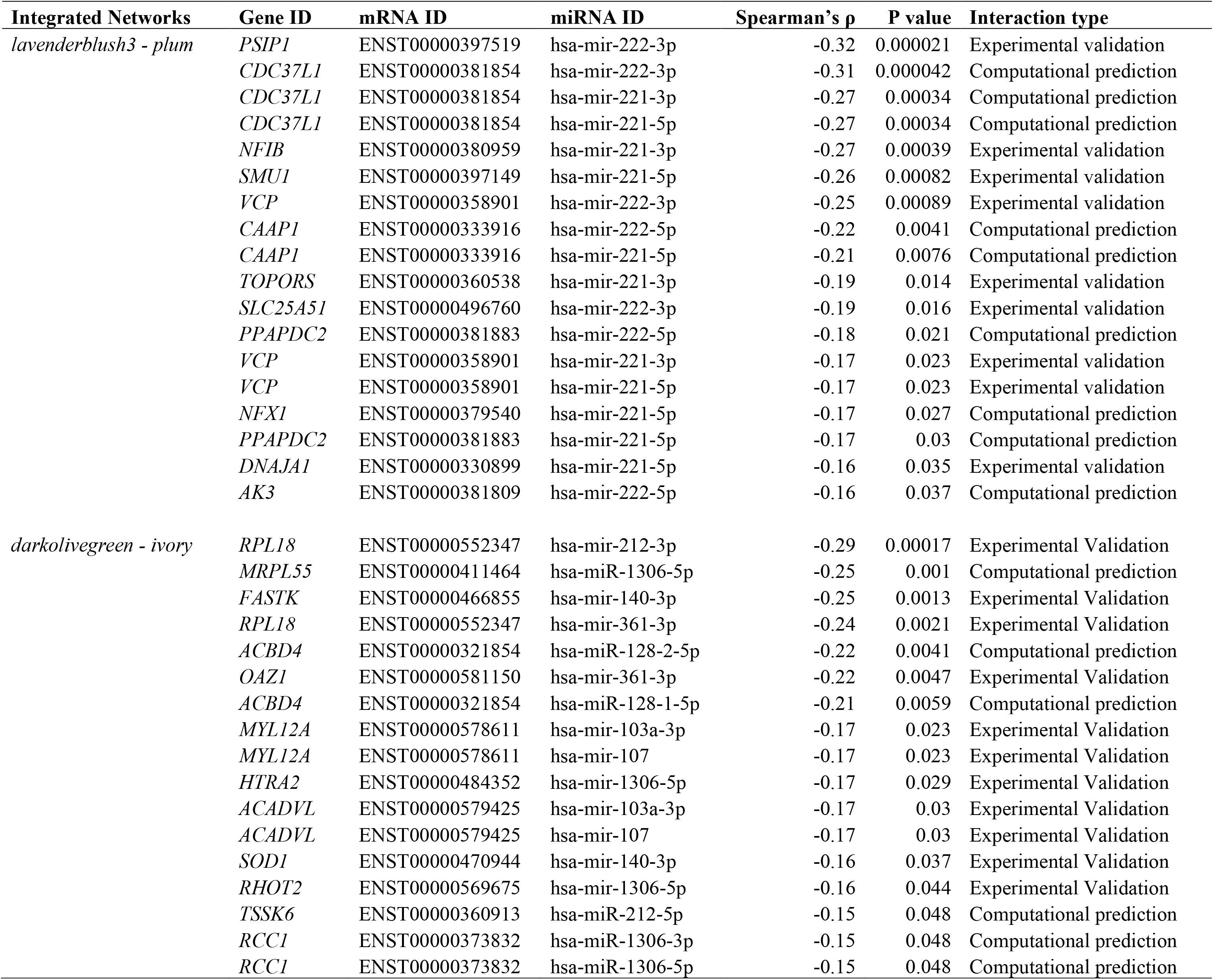
Predicted and validated mRNA - miRNA network interactions.

**Figure 3.**
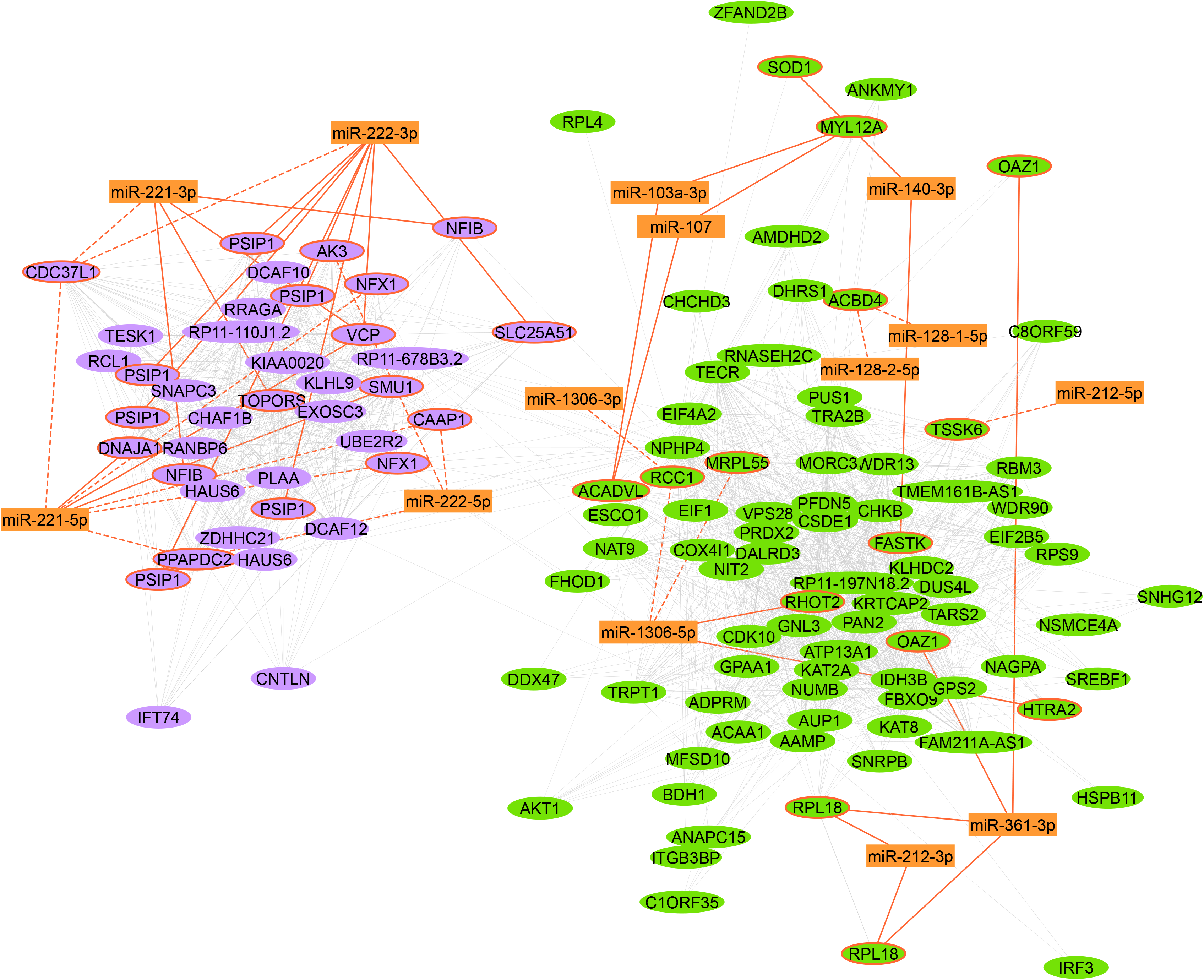
Integration of mRNA - miRNA networks. The mRNA transcripts (oval nodes) from the *lavenderblush3* and *darkolivegreen* modules are arranged based on co-expression similarity (grey edges). Four common miRNA isoforms (rectangular nodes) were validated (orange solid edges) or predicted (orange dashed edges) to regulate the expression of genes in the *lavenderblush3* network module, while 10 miRNA isoforms were validated or predicted to regulate genes in the *darkolivegreen* module. Genes targets of miRNAs indicated by orange border.

Replication analysis of the mRNA results used RNA microarray data from the AOCS cohort. In total, 140 microarray probes in the AOCS dataset (73.68%) matched to differentially expressed genes from TCGA-HGSOC, 28 (20%) of which replicated with the same direction of effect (**Table S6**). In addition, the *lavenderblush3* and *darkolivegreen* mRNA network modules replicated in the AOCS cohort (p < 0.0001) (**Figure 4A, B**).

**Figure 4.**
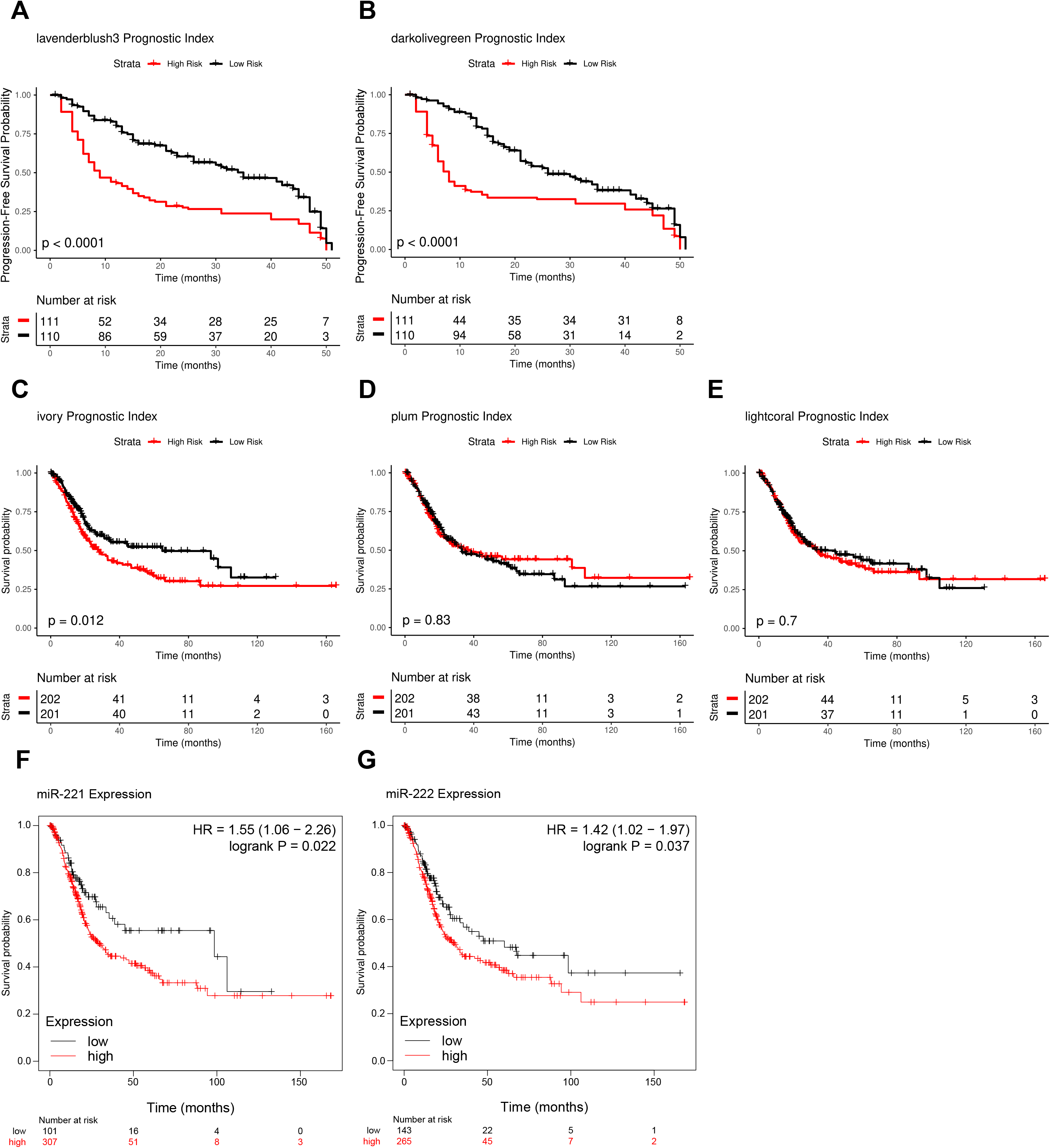
Replication analysis of the gene co-expression networks. (A-B) Prognostic Index Kaplan-Meier analysis of patient PFS association with the prognostic index of mRNA networks *lavenderblush3* (A) and *darkolivegreen* (B). (C-E) Prognostic Index Kaplan-Meier analysis of patient OS association with the prognostic index of miRNA networks *ivory* (C), *plum* (D), and *lightcoral* (E). (F-G) Kaplan-Meier analysis of patient OS with the expression of miR-221 (F) and miR-222 (G). PFS, progression-free survival; OS, overall survival.

Replication of the miRNA findings used miRNA-seq data from the BLCA cohort of TCGA. Of the 16 differentially expressed miRNAs isoforms in the HGSOC data, 13 (81.25%) were available in the TCGA-BLCA data and 7 (53.85%) replicated with the same direction of effect (**Table S7**). In addition, the *ivory* miRNA network module replicated in the TCGA-BLCA cohort (p = 0.012), while the *lightcoral* and *plum* modules did not replicate (**Figure 4C-E**). However, differential expression of miR-221 and miR-222, which make up the *plum* network, replicated in this cohort using Kaplan-Meier analysis (p = 0.022 and p = 0.037) (**Figure 4F, G**).

## 4. Discussion

In this study, we analyzed mRNA and miRNA sequencing data from chemotherapy-naïve tumors of HGSOC patients to identify transcripts and networks associated with chemotherapy response. Our findings implicate novel and known biological pathways that replicated in independent cancer cohorts. In addition, we identified potential interactions among miRNAs and mRNAs, as well as eQTLs that potentially regulate the associated transcripts. Thus, our results provide an integrative, multi-omics view of biological networks associated with chemotherapy response.

We identified one mRNA co-expression network (*lavenderblush3*) significantly upregulated in platinum sensitive patients, which replicated in the AOCS. This module consists of genes involved in ubiquitin-mediated proteolysis in the endoplasmic reticulum (ER). We also detected a significant downregulation of genes responsible for translation initiation in sensitive patients. These findings suggest that the unfolded protein response (UPR), a cellular process responsible for resolving ER stress, may be increasingly activated in sensitive patients compared to resistant cases. The UPR alleviates ER stress through several pathways, including increased ER-associated protein degradation (ERAD) to remove misfolded proteins, and inhibition of translation to reduce protein load in the ER [17]. ER stress promotes cisplatin resistance in OC cell lines [18] and the upregulation of ERAD genes such as *VCP* in the *lavenderblush3* module is associated with longer OS and platinum sensitivity in HGSOC cohorts [9, 19]. Eighteen of 31 module genes were also reported in an earlier study of microarray data from the same TCGA-HGSOC cohort [9].

We identified a second mRNA co-expression network associated with chemotherapy resistance in our HGSOC cohort that replicated in the AOCS. The *darkolivegreen* module included genes associated with fatty acid metabolism (*SREBF1*, *ACAA1*, *ACADV*L), and the protein kinase B oncogene (*AKT1*), which promotes de-novo lipid biosynthesis in cancer [20]. SREBF1 is a key enzyme for cholesterol and fatty acid synthesis, and an essential gene for OC tumor growth [21]. Specifically, *SREBF1* is activated by *AKT1*, promoting fatty acid synthesis [22], which favors cell proliferation in OC [23]. Expression of *ACADVL*, involved in the β-oxidation of long-chain fatty acids, is linked to OC metastasis and cell survival [24]. Our findings indicate the upregulation of these lipid metabolism genes among chemotherapy resistant patients. Lipid metabolism dysregulation activates the UPR, which triggers lipid metabolism-based adaptations in the cell through several pathways, including *SREBF1* regulation [25]. The interaction of these pathways may present a link between our two gene co-expression modules and warrants further study.

Differential expression analysis identified a downregulation of mRNA transcripts involved in the adaptive immune system, which is associated with chemoresistance. Previous studies reported that a high tumor immune score is a strong predictor of chemosensitivity in HGSOC [26]. In addition, there are potential links between this immune response activation, UPR and lipid metabolism. ER stress can induce pro-inflammatory cytokine production and UPR activation in tumor cells [27], which can disrupt dendritic cell function in the OC tumor microenvironment [28]. Moreover, dendritic cell function can also be inhibited by increased lipid uptake in various cancers [29].

The *ivory* miRNA co-expression network module, associated with chemotherapy sensitivity in HGSOC and BLCA, is involved in the negative regulation of lipid transport. This enrichment is mainly mediated by miR-128-1 and miR-128-2, which play a key role in cholesterol and lipid homeostasis through their suppression of the ABCA1 cholesterol efflux transporter and the low-density lipoprotein receptor (LDLR) [30, 31]. MiR-148a is also a regulator of these key genes [30], which is significantly downregulated in resistant patients. The overexpression of *ABCA1* is associated with reduced survival in OC patients [32], and levels of LDLR are increased in chemoresistant OC cell lines [33]. In addition, overexpression of miR-128 promotes sensitivity to cisplatin in previously resistant OC cells [34]. Our results are consistent with the chemosensitivity-promoting role of miR-128 and its potential activity in cholesterol efflux inhibition alongside miR-148a in this cohort.

The *plum* miRNA co-expression network consists of miR-221 and miR-222 isoforms, which have been implicated in the development of chemotherapy resistance in OC. Expression of miR-221/miR-222 transcripts is high in cisplatin-resistant OC cell lines, and their inhibition increases cellular sensitivity [35]. Over-expression of miR-221 and miR-222 promotes proliferation of OC cell lines [36, 37] and reduced disease-free and overall survival [36]. Thus, our findings are consistent with earlier studies showing increased activity of miR-221 and miR-222 in chemoresistant tumors.

Integrative analysis of mRNA-seq and miRNA-seq datasets identified potential interactions of the associated transcript co-expression modules. The overexpression of miR-221/222 in resistant patients may be inhibiting the chemosensitivity-associated *lavenderblush3* mRNA network, revealing a novel potential mechanism of chemotherapy resistance. This finding, combined with the accumulating evidence of miR-221/miR-222 involvement in chemoresistance, may point to a promising avenue for therapeutic intervention. However, overexpression of miR-221/miR-222 promotes UPR-induced apoptosis in hepatocellular carcinoma (HCC) cells [38]. Additionally, ER stress suppresses miR-221/miR-222 in HCC, promoting resistance to apoptosis. The contribution of this mechanism to chemotherapy response in HGSOC is currently unclear and presents an area for future investigation.

We also identified potential regulation of the *darkolivegreen* mRNA module by the *ivory* miRNA network, which may inhibit lipid metabolism in chemotherapy sensitive patients. As increased lipid metabolism by cancer cells is a known mechanism of chemoresistance in HGSOC, this miRNA-mediated inhibition may present a novel mechanism of chemotherapy sensitivity.

Finally, cis-eQTL analysis identified known and novel genomic variants correlated with the expression of mRNAs and miRNAs, which are associated with lipid-related phenotypes. High HDL and triglyceride levels have been correlated with increased cancer stage at diagnosis in OC patients [39]. In addition, advanced-stage OC patients with high LDL levels have a shorter PFS than patients with normal levels [40]. Further investigation of these eQTLs is necessary to further elucidate their role in platinum-based chemotherapy resistance and HGSOC prognosis.

## 5. Conclusion

Our study provides novel insight of the underlying mechanisms modulating resistance to platinum-based chemotherapy in HGSOC. Specifically, we conducted whole-transcriptome analysis of mRNA-seq and miRNA-seq data to generate novel mechanistic hypotheses using both univariate and network methods. Moreover, we integrated this data with miRNA-seq and genome-wide SNPs to determine potential regulation of the associated transcripts and networks. Our findings implicate novel and known signaling pathways and networks associated with chemotherapy response in HGSOC as well as regulators, which could become novel drug targets. Further studies are needed to validate these findings in other cancers.

## Abbreviations

AOCS: Australian Ovarian Cancer Study
BLCA: Bladder Urothelial Carcinoma
eQTL: Expression Quantitative Trait Locus
HGSOC: High-Grade Serous Ovarian Cancer
OC: Ovarian Cancer
OS: Overall Survival
PFS: Progression-Free Survival
TCGA: The Cancer Genome Atlas
WGCNA: Weighted Gene Co-Expression Network Analysis

## CRediT authorship contribution statement

**Danai Georgia Topouza:** Investigation, Formal analysis, Software, Visualization, Validation, Data curation, Writing – original draft, Writing – review & editing. **Jihoon Choi:** Formal analysis, Software, Data curation, Writing – review & editing. **Sean Nesdoly:** Software, Data curation. **Anastasiya Tarnouskaya:** Data curation. **Christopher J.B. Nicol:** Writing – review & editing. **Qing Ling Duan:** Conceptualization, Supervision, Methodology, Resources, Funding acquisition, Writing – review & editing.

## Acknowledgements

Computations in this manuscript were performed on resources and with support provided by the Centre for Advanced Computing (CAC) at Queen’s University in Kingston, Ontario. The CAC is funded by the Canada Foundation for Innovation, the Government of Ontario, and Queen’s University.

## Funding sources

D.G.T. is funded by Queen’s University and internal awards in the Department of Biomedical and Molecular Sciences. Q.L.D. receives funding from the Canadian Institutes of Health Research and Queen’s University National Scholar Award. The funders did not play any role in the study design, data collection, data analysis, data interpretation, writing of the manuscript or decision to publish results.

## Data statement

Transcriptomics, genomics, and clinical data used for the analysis of the TCGA-OV and TCGA-BLCA cohorts can be accessed and downloaded from the Genomic Data Commons (GDC) Data Portal (https://portal.gdc.cancer.gov/). Gene expression and clinical data of the AOCS cohort can be accessed and downloaded from the Gene Expression Omnibus (GEO) database (https://www.ncbi.nlm.nih.gov/geo/query/acc.cgi?acc=GSE9899).

## Ethics declarations

Controlled access to datasets from The Cancer Genome Atlas (TCGA) was provided by the National Institute of Health (NIH). The terms of access are outlined in the Data Use Certification Agreement: https://dbgap.ncbi.nlm.nih.gov/aa/wga.cgi?view_pdf&wlid=10654&tlsid=274. Details of the Human subject protection and data access policies implemented by TCGA are as described: https://www.cancer.gov/about-nci/organization/ccg/research/structural-genomics/tcga/history/policies/tcga-human-subjects-data-policies.pdf. Local ethics clearance for human subject research was granted by Queen’s University.

## Declaration of interest statement

None declared.

## Supporting Information

### Supplementary Methods

**Supplementary Data 1.** mRNA-microRNA interaction annotations.

**Table S1.** mRNA differential expression analysis results.

**Table S2.** microRNA differential expression analysis results.

**Table S3.** Significant WGCNA module mRNAs and correlation to chemotherapy resistance.

**Table S4.** Significant WGCNA module microRNAs and correlation to chemotherapy resistance.

**Table S5.** eQTL analysis results and annotation.

**Table S6.** Kaplan-Meier validation results for differentially expressed mRNAs.

**Table S7.** Kaplan-Meier validation results for differentially expressed microRNAs.

## Notes

### Competing Interest Statement

The authors have declared no competing interest.

### Summary of Updates

Reformatted manuscript and reduced word count; Supplemental files updated.

