## Supplementary Methods for "Novel regulatory and transcriptional networks associated with resistance to platinum-based chemotherapy in ovarian cancer"

*1. Chemotherapy response classification*

Sensitive patients remained cancer-free one year after completion of chemotherapy, whereas resistant patients experienced cancer recurrence within 6 months of chemotherapy completion. We excluded patients who developed new tumors between six months to one year from chemotherapy completion to enrich for genetic differences between the drug response groups. Included patients received platinum-based adjuvant chemotherapy, which consisted of a platinum agent (3.66 – 3.90% of patients) or a combination of a platinum agent and a taxane (96.10 – 96.34% of patients) (**Table 1**). We excluded patients who were treated with an alternative therapy instead of platinum-based adjuvant chemotherapy, patients without a tumor recurrence who passed away within six months of chemotherapy completion, patients who were followed for less than one year after chemotherapy completion, and patients without transcriptomic data.

*2. Study design*

An analysis pipeline for mRNA-sequencing data was curated as summarized in **Figure S1** and detailed in the following sections. Our pipeline applied select software and tools for quality control of raw sequence reads, alignment to a reference genome, quantification of reads into transcript isoform expression values, and univariate and multivariate analysis for correlation of transcripts with chemotherapy response.

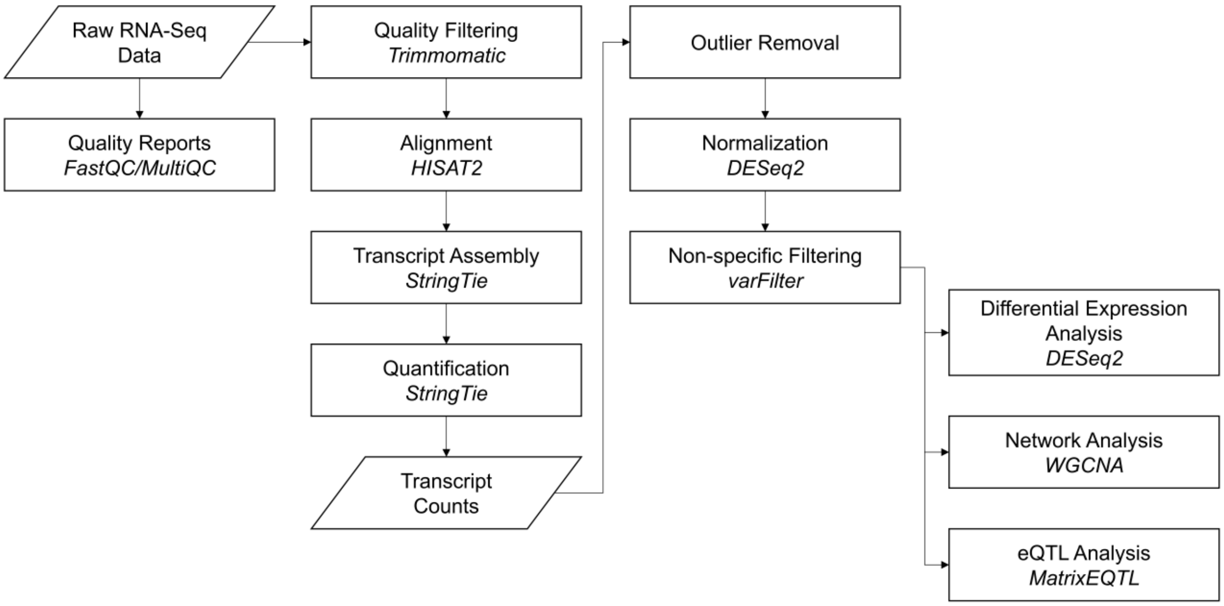


**Figure S1. Flowchart of RNA-seq analysis pipeline.** First, reports of read sequence quality were generated using FastQC [1] and MultiQC [2], followed by trimming of low quality RNA-seq reads with Trimmomatic [3]. Filtered reads were then aligned to the hg19 human reference genome [4] using HISAT2 (hierarchical indexing for spliced alignment of transcripts) [5], and quantified by StringTie [6]. Next, transcript expression was normalized using the DESeq2 R package [7], and highly variable transcripts were selected with the varFilter function of the genefilter R package [8]. Finally, transcript expression was used to test for differential expresssion using DESeq2 and to construct co-expression networks using the WGCNA (weighted gene co-expression network analysis) R package [9]. Moreover, the sequence data was integrated with genomics data from the same patients to determine eQTLs (expression quantitative trait loci) using the MatrixEQTL R package [10].

*3. RNA-Sequencing data preprocessing*

Raw mRNA-sequencing reads from frozen chemotherapy-naïve tumor samples of high-grade serous ovarian cancer (HGSOC) patients were obtained from the TCGA database [11] on August 31st, 2017 using the TCGAbiolinks R package [12]. Patient mRNA-seq data were filtered for base quality using the Trimmomatic software package [3]. Bases were evaluated in a window of 4 bases progressing from the 5' end of each read to the 3' end. Each read was trimmed when the average quality within the window dropped below 15 on the PHRED base 33 quality scale, which indicates a 3.1% probability of error [13]. Sequences matching Illumina HiSeq adapters and Illmina PCR primers were removed.

Patient RNA-sequence reads were aligned to the hg19 reference genome (GATK resource bundle) [14] using the HISAT2 software [5], with a mean overall alignment rate of 96.38%. The aligned reads were assembled into transcripts by the StringTie software package [6] and quantified using the GRCh37.87 transcript list defined by Ensembl (release 92) [15]. This procedure resulted in the quantification of gene transcripts and isoforms (n=196,464) for each patient.

Aligned and quantified frozen chemotherapy-naïve tumor microRNA-sequencing data (Level 3) for the TCGA HGSOC cohort were obtained from the Broad Institute Genome Data Analysis Center Firehose [16] (http://gdac.broadinstitute.org) on September 17th, 2018 (TCGA data version 2016_01_28 for OV). Patient data were acquired in the form of quantified microRNA-seq isoform counts (n=29,382) in the .txt format. The microRNA (miRNA) isoform data had been aligned to GRCh36/hg18, and quantified using TCGA's modified version of the British Columbia Genome Sciences Centre miRNA profiling pipeline [17].

*4. Patient-level quality control*

Patient mRNA data were iteratively filtered for outliers based on Spearman's correlation of gene expression between patients, using Tukey's outlier labelling threshold ($Q1 - \alpha * IQR$) [18] and fine-tuning the α value based on our sample size ($\alpha = 2.3$) [19] as described by Panarelli *et al.* [20]. This process removed 4 patients. Patient miRNA expression profiles were also filtered for outliers using the above method, with an α value of 2.4 based on the larger number of patients in the miRNA cohort. This process removed 9 patients (**Figure S2**). Patient age at diagnosis was found to be significantly higher in resistant patients (**Table 1**) and was used as a covariate in our analyses.

*5. Transcript-level quality control*

Transcript count data were normalized using the median of ratios method from the DESeq2 R software package [7]. Next, we applied non-specific filtering to the mRNA expression dataset using median absolute deviation (MAD) [8], and retained the top 25% of mRNA transcripts with the highest variance among patients for further analysis (n=49,116). As for non-specific filtering of the miRNA dataset, we omitted isoforms with variance below the median value, resulting in 14,691 remaining miRNA transcripts. In addition, we removed miRNA isoforms with low expression among patients (isoform count mean < 1), resulting in 4,479 miRNAs for further analysis.

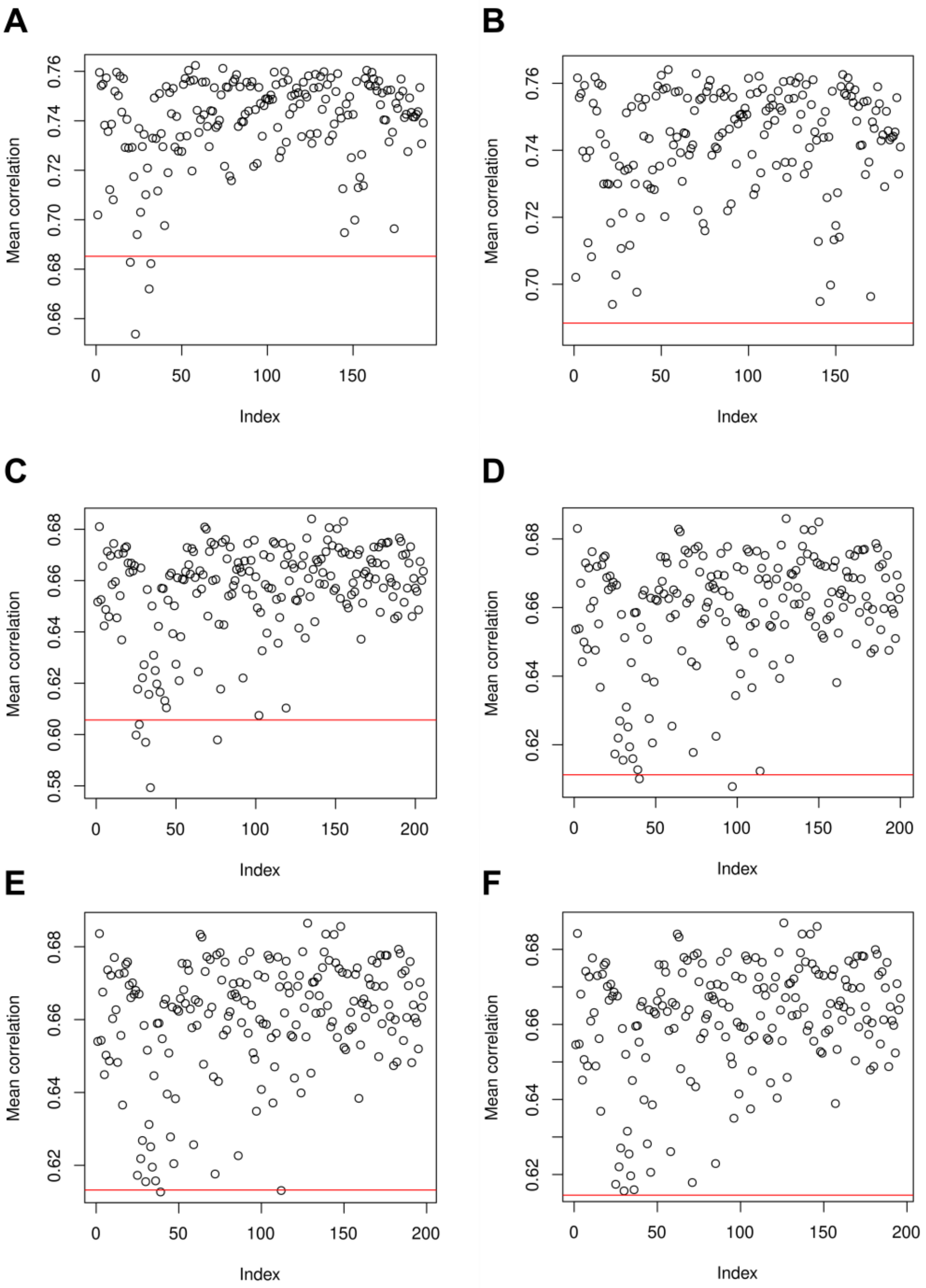


**Figure S2. Outlier detection.** (A, B) Scatterplot of mean Spearman correlation for mRNA expression profiles. Patients who fell below the threshold for minimum correlation (red line), as computed by (Q1 – 2.3 * IQR), were removed (A). The correlations and threshold were then calculated again, until no subjects fell below the correlation threshold (B). (C-F) Scatterplot of mean Spearman correlation for miRNA expression profiles. Patients who fell below the minimum correlation threshold (red line), as computed by (Q1 – 2.4 * IQR), were removed. Patient correlation and threshold were then re-calculated four times, until no subjects fell below the correlation threshold (F).

*6. Differential expression analysis*

Volcano plot figures were generated using the R package ggplot2 (v. 3.3.2) [21].

*7. Pathway enrichment analysis*

Three differentially expressed miRNAs (miR-151a, miR-103a-1, miR-203a) were re-annotated with the newest miRbase [22] identifiers before pathway analysis with g:Profiler. Three of the 16 miRNAs (miR-320a, miR-1974, and miR-886) are not yet identified in the g:Profiler database and not included in the pathway enrichment analysis.

*8. Weighted transcript co-expression network analysis*

Patient transcript expression data were analyzed using the weighted gene co-expression network analysis (WGCNA) R package [9]. First, this tool calculated the Pearson correlation using the expression values of all transcripts. Next, an adjacency function was used to transform each correlation into a connection strength, forming a network where each node is a gene, and each connecting edge corresponds to the strength of the correlation between nodes. The power adjacency function raised the correlations to a power $\beta$, making strong connections stronger and weak connections weaker. This step made the connectivity distribution of our gene expression network approach that of a scale-free network. Fine tuning indicated $\beta = 10$ to be optimal for the mRNA dataset, and $\beta = 9$ to be optimal for the miRNA dataset (**Figure S3**).

In addition to adjacency, WGCNA used the topological overlap of two nodes, which is the number of neighbor nodes they share, to form gene clusters. A Topological Overlap Matrix (TOM) was constructed, where the topological overlap of two transcripts is a function of their adjacency and number of shared connections. The TOM-based dissimilarity was then used in hierarchical clustering to generate gene modules. We used a minimum module size of 30 to encourage larger gene clusters, as recommended by the WGCNA manual [23].

After clustering, we merged modules that were not sufficiently distinct from each other to produce robust networks that better capture biological pathways. We calculated a module membership value for each transcript that shows how well a transcript fits into each of the available modules. 50% of transcripts from each module were tested for membership. If more than 25% of transcripts tested have a higher membership value for a module other than their own, those two modules were merged. This process was performed in rounds until no more modules could be merged.

Finally, principal component analysis was conducted for all transcripts within a module, resulting in a value called an eigengene. We used these eigengene values and the patient age covariate to construct a generalized linear model that revealed modules with significant correlation with chemotherapy response. Networks with a significant association were functionally annotated using pathway analysis. Cytoscape was used for network visualisation [24].

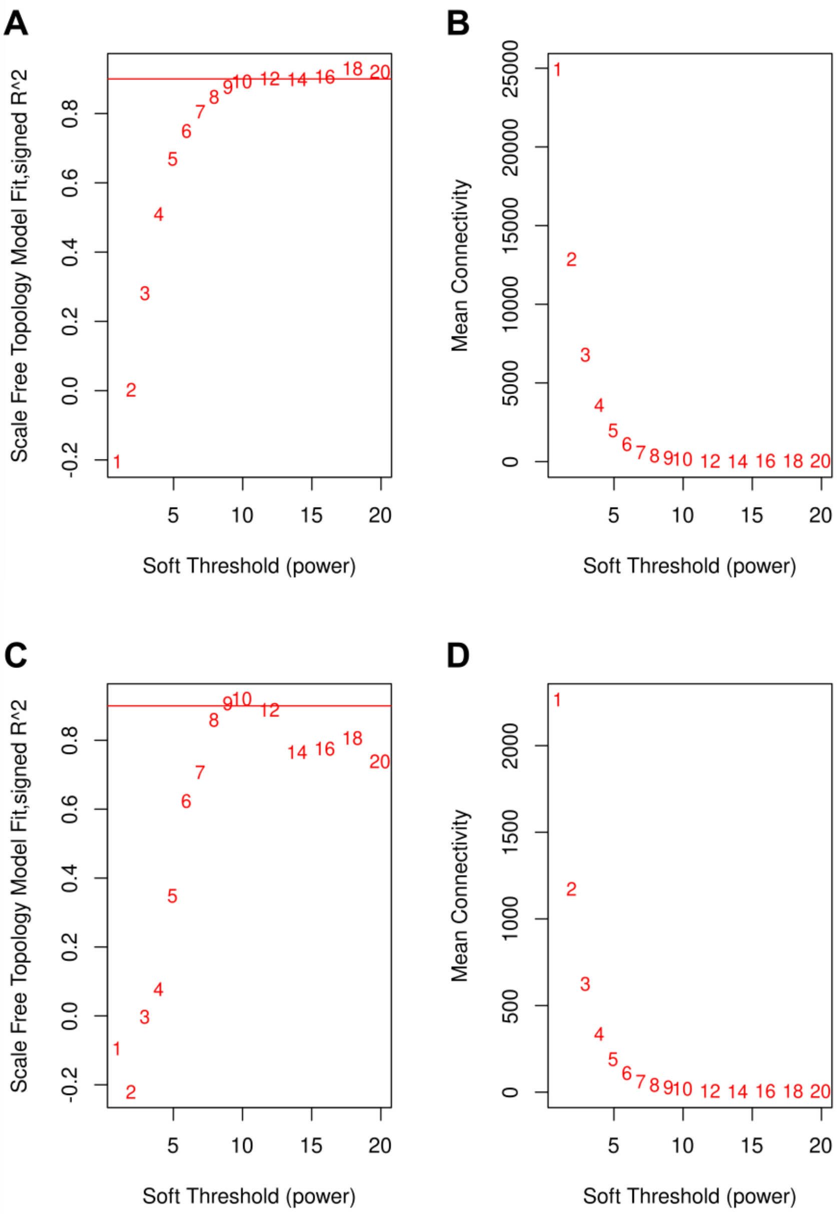


**Figure S3. Selection of soft thresholding power for network analysis.** (A,C) Transcript network connectivity fit to the scale-free topology as a function of the power parameter β. Our transcript expression network connectivity had a 90% similarity to the scale-free topology at β = 10 for mRNA (A) and at β = 9 for miRNA (C) data. (B,D) Mean node connectivity as a function of β for mRNA (B) and miRNA (D) data. As the network fit to the scale-free topology increases, only a few nodes retain high connectivity (hubs), while the majority of nodes have very low connectivity. Therefore, as $\beta$ increases, the mean network connectivity decreases.

*9. Expression Quantitative Trait locus (eQTL) analysis*

Germline single nucleotide polymorphism (SNP) data were profiled in normal tissues of HGSOC patients in the TCGA cohort using the Affymetrix SNP Array 6.0. 47,960,330 genetic polymorphisms for 262 patients were obtained after phasing and imputing as described by Choi *et al.* [25]. The imputed data was then processed for quality control using PLINK 1.9 [26]. We first performed patient-level quality control: 20 patients were removed due to low ($F < -0.05$) or excessive ($F > 0.05$) heterozygosity, and 2 patients were removed due to high genetic relatedness (pi-hat > 0.9), leaving 240 patients for further analysis. We then performed variant-level quality control, starting with linkage disequilibrium (LD)-based variant pruning. This process evaluated variants in a window of 50 SNPs, which shifted by 5 SNPs after every iteration, and removed variants with a variance inflation factor (VIF) larger than 2 ($r^2 > 0.5$). 7,075,175 independent SNPs were retained after LD pruning. After checking for allelic independence with the Hardy-Weinberg equilibrium, 112 more variants were removed. Finally, we retained SNPs with a minor allele frequency ≥ 5% and variant missingness < 10%, resulting in 1,722,608 variants to be used for further analysis. 167 patients from the RNA-Seq cohort and 178 patients from the miRNA-Seq cohort had genomic data available following quality control.

Next, we tested the association of common patient polymorphisms (n=1,722,608) with mRNA transcript expression (n=196,464) or miRNA isoform expression (n=29,382), in order to determine novel expression quantitative trait loci (eQTLs) in our dataset. We used the MatrixEQTL R package [10] to test SNPs located within 1 Mb of recorded transcripts [27] (cis-eQTLs) for association with transcript expression. Based on this distance, 1,676,365 common SNPs were tested for association with mRNA expression, while 356,189 common SNPs were tested for association with miRNA expression. miRNA isoform locations were converted to hg19 coordinates using UCSC's liftOver utility [28].

We identified 121,900 significantly associated SNP-mRNA transcript pairs and 36,995 significantly associated SNP-miRNA isoform pairs after false discovery rate correction. Of those, 248 SNPs were associated with 55 mRNA transcripts found to be significant in differential expression or network analysis, while 20 SNPs were associated with 7 miRNAs that were significant in differential expression or network analysis (**Table S5**). We mapped these SNPs to an rsID using the dbSNP Build150 Human Variation Set [29], and functionally annotated them using HaploReg v4.1 [30].

*10. Replication analysis*

The replication of our differential expression analysis findings was performed by evaluating the association of each transcript with progression-free survival (PFS) in an independent ovarian cancer cohort using KM-Plotter [31]. The Kaplan-Meier analysis for the replication of mRNA results was performed on microarray-collected tumor expression data from a cohort of 285 patients from the Australian Ovarian Cancer Study (AOCS) (GSE9891) [32]. We replicated our miRNA results by using the KM-Plotter tool to test for the association of miRNA expression and overall survival (OS) in an independent cancer cohort. No suitable independent large ovarian cancer miRNA dataset was available to be used for validation; the TCGA bladder urothelial carcinoma (TCGA-BLCA) cohort (n=409) [33] was chosen, as bladder cancer is also typically treated with platinum-based combination chemotherapy [34]. OS was used as the target phenotype, as PFS was not available for this cohort.

The replication of our network analysis results was performed using the Prognostic Index (PI) estimation method as described by Aguirre-Gamboa *et al.* in the SurvExpress biomarker validation tool [35]. We used the AOCS and TCGA-BLCA datasets to validate our mRNA and miRNA networks respectively. The AOCS expression microarray data were retrieved from GEO and processed as described by Choi *et al.* [25] before analysis, while the normalized TCGA-BLCA data (Level 3) were retrieved from GDC Firehose [16]. Figures 4A-E were generated using the R package survminer (v. 0.4.8) [36]. Figures 4F,G were generated by the KM Plotter tool.
